# Growth differentiation factor 15 increases in both cerebrospinal fluid and serum during pregnancy

**DOI:** 10.1101/2021.03.10.434730

**Authors:** Ulrika Andersson-Hall, Pernilla Svedin, Carina Mallard, Kaj Blennow, Henrik Zetterberg, Agneta Holmäng

## Abstract

**Aim:** Growth differentiation factor 15 (GDF15) increases in serum during pregnancy to levels not seen in any other physiological state and is suggested to be involved in pregnancy-induced nausea, weight regulation and glucose metabolism. The main action of GDF15 is regulated through a receptor of the brainstem, i.e., through exposure of GDF15 in both blood and cerebrospinal fluid (CSF). The aim of the current study was to measure GDF15 in both CSF and serum during pregnancy, and to compare it longitudinally to non-pregnant levels.

**Methods:** Women were sampled at elective caesarean section (n=45, BMI=28.1±5.0) and were followed up 5 years after pregnancy (n=25). GDF15, insulin and leptin were measured in CSF and serum. In addition, glucose, adiponectin and Hs-CRP were measured in blood.

**Results:** GDF15 levels were higher during pregnancy compared with follow-up in both CSF (385±128 vs. 115±32 ng/l, p<0.001) and serum (73789±29198 vs. 404±102 ng/l, p<0.001). CSF levels correlated with serum levels during pregnancy (p<0.001), but not in the non-pregnant state (p=0.98). Both CSF and serum GDF15 were highest in women carrying a female fetus (p<0.001), previously linked to pregnancy-induced nausea. Serum GDF15 correlated with the homeostatic model assessment for beta-cell function and placental weight, and CSF GDF15 correlated inversely with CSF insulin levels.

**Conclusion:** This, the first study to measure CSF GDF15 during pregnancy, demonstrated increased GDF15 levels in both serum and CSF during pregnancy. The results suggest that effects of GDF15 during pregnancy can be mediated by increases in both CSF and serum levels.

## Introduction

Growth differentiation factor 15 (GDF15), a member of the transforming growth factor-beta family, was first discovered to be involved in inflammation and stress pathways but has also emerged as a potentially important metabolic regulator. (1–3) GDF15 has for example been shown to induce weight loss (probably through appetite suppression and decreased food intake), affect energy expenditure and motivation to exercise, and improve glucose tolerance. (4–9)

Research interest has recently grown regarding GDF15 during pregnancy as substantial and progressive increases in serum GDF15 have been shown from early to late pregnancy, ending up with serum levels much higher than in any other physiological or pathophysiological state. (10–12) Pregnancy is marked by major metabolic and physiological changes, such as increases in appetite, body weight, insulin resistance and inflammation. (13, 14) GDF15 may play an important role in all these areas, and has been found during pregnancy to be linked to altered glucose metabolism, (12, 15) and pregnancy-induced nausea. (16–18)

Food intake and energy expenditure are primarily controlled by the central nervous system, with the arcuate nucleus of the hypothalamus identified as a key area. Several appetite-suppressing/stimulating neuropeptides have been shown to change both in the circulation and in the cerebrospinal fluid (CSF) during pregnancy. (19, 20) GDF15 signals through the glial cell line-derived neurotrophic factor (GDNF) family receptor alpha-like (GFRAL), a receptor believed to be present only in the area postrema (AP) and the nucleus of the solitary tract (NTS) regions of the brainstem,(21, 22) which in turn signals to the arcuate nucleus in the hypothalamus. The AP/NTS region is also generally believed to be the main signaling center for nausea.(23) AP has a highly permeable blood brain barrier (BBB) compared to other brain regions, and can therefore receive signals both from the blood and CSF, whereas NTS is separated from AP with a more solid BBB.(24)

Local hypothalamic expression of GDF15 or intracerebroventricular injections of recombinant human GDF15 in mice resulted in a direct central action which induced anorexia and weight loss.(25) Although numerous recent human studies have measured GDF15 in the circulation, few have examined GDF15 concentrations in the CSF. Studies to date have shown approximately 50% increased GDF15 levels in CSF of patients with neurodegenerative disorders or glioblastomas.(26–28) However, the effect of pregnancy on CSF levels - a physiological state with up to 200-fold increases in circulating GDF15 – is not known. The aim of this study was to determine the concentration of GDF15 in both CSF and serum during pregnancy and to establish whether CSF GDF15 was different in the pregnant compared to the non-pregnant state in a cohort of women sampled at elective cesarean section and after pregnancy.

## Materials and Methods

### Ethical approval

The study was approved by the ethical committee at the University of Gothenburg (dnr 402-08/dnr 750-15). Informed consent was obtained from all participants.

### Study cohort

Women were recruited at admission for elective cesarean section as previously described,(19) and followed up 5 years after pregnancy.(29) All women with serum and/or CSF samples available from the previous study were included in the present study. Out of 74 women in the original study, 45 women had remaining samples from the cesarean section and 24 women had samples from the 5-year follow-up. Inclusion criteria in the original study were uncomplicated pregnancy and good health, judged from the medical history. At entry, all subjects were normoglycemic, nonsmokers, and did not consume alcohol. Dieting and use of weight-loss supplements within 6 months before pregnancy were excluding criteria. The characteristics and blood/CSF measurements for the women in the study are presented in Table 1. A drop-out analysis showed that women who did not attend the follow-up had higher pre-pregnancy BMI, and pregnancy glucose and insulin compared with women attending both visits (BMI, 30 ± 4 vs 27 ± 4 kg/m^2^, *p* = 0.01; glucose, 4.5 ± 1.3 vs 3.9 ± 0.6 mmol/l, *p* = 0.04; insulin, 13.9 ± 8.0 vs9.0 ± 5.4 mU/l, *p* = 0.02). There were no differences in age, gestational weight gain, placental weight or birth weight (*p* > 0.3), and importantly, there was no difference in GDF15 levels in serum or CSF at pregnancy between women that attended one or two visits (*p* > 0.5).

**Table 1.**
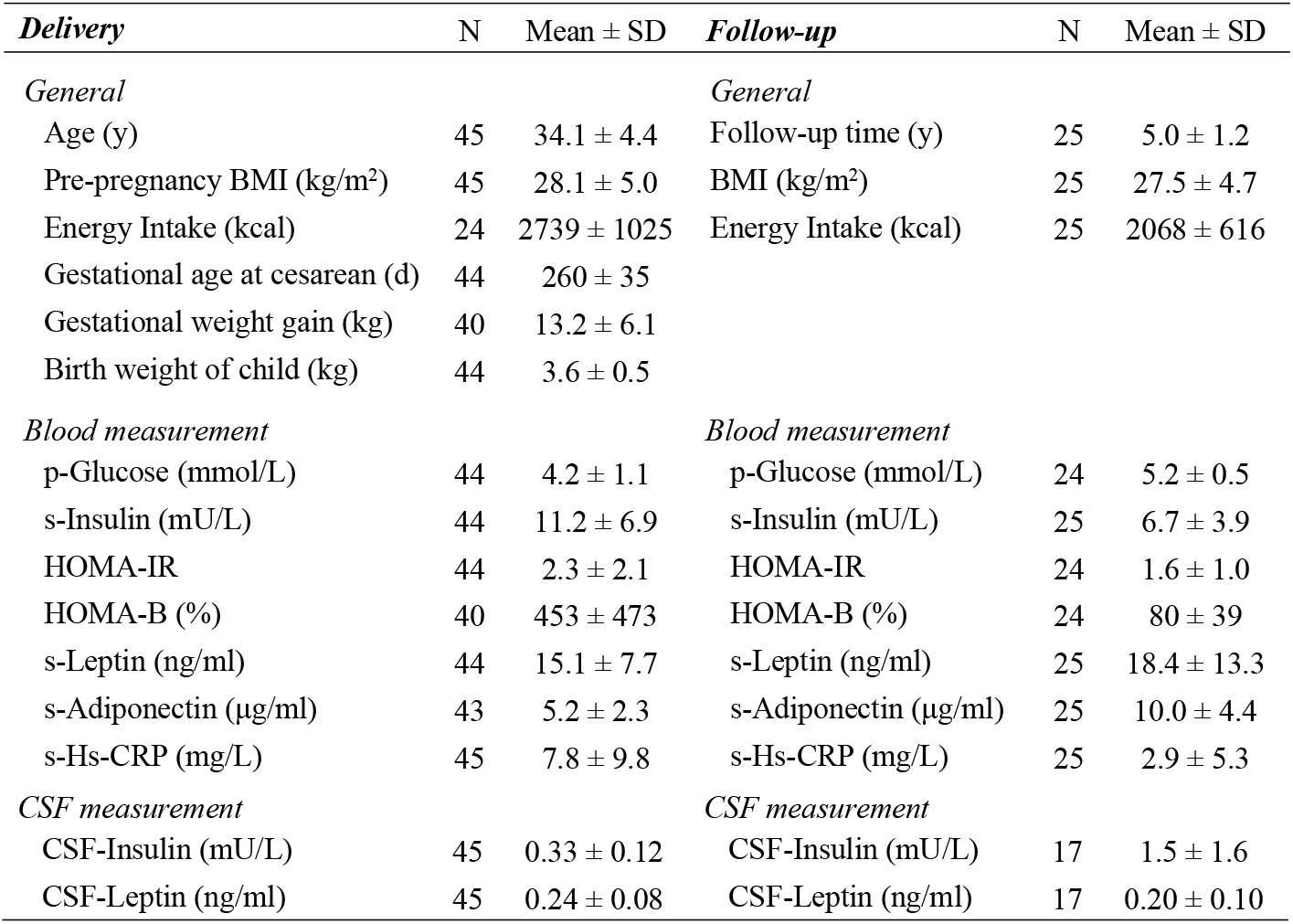
Maternal characteristics at delivery and at follow-up

| ***Delivery*** | N | Mean ± SD | ***Follow-up*** | N | Mean ± SD |
| --- | --- | --- | --- | --- | --- |
| *General* |  |  | *General* |  |  |
| Age (y) | 45 | $34.1 \pm 4.4$ | Follow-up time (y) | 25 | $5.0 \pm 1.2$ |
| Pre-pregnancy BMI (kg/m<sup>2</sup>) | 45 | $28.1 \pm 5.0$ | BMI (kg/m<sup>2</sup>) | 25 | $27.5 \pm 4.7$ |
| Energy Intake (kcal) | 24 | $2739 \pm 1025$ | Energy Intake (kcal) | 25 | $2068 \pm 616$ |
| Gestational age at cesarean (d) | 44 | $260 \pm 35$ | | | |
| Gestational weight gain (kg) | 40 | $13.2 \pm 6.1$ | | | |
| Birth weight of child (kg) | 44 | $3.6 \pm 0.5$ | | | |
| *Blood measurement* |  |  | *Blood measurement* |  |  |
| p-Glucose (mmol/L) | 44 | $4.2 \pm 1.1$ | p-Glucose (mmol/L) | 24 | $5.2 \pm 0.5$ |
| s-Insulin (mU/L) | 44 | $11.2 \pm 6.9$ | s-Insulin (mU/L) | 25 | $6.7 \pm 3.9$ |
| HOMA-IR | 44 | $2.3 \pm 2.1$ | HOMA-IR | 24 | $1.6 \pm 1.0$ |
| HOMA-B (%) | 40 | $453 \pm 473$ | HOMA-B (%) | 24 | $80 \pm 39$ |
| s-Leptin (ng/ml) | 44 | $15.1 \pm 7.7$ | s-Leptin (ng/ml) | 25 | $18.4 \pm 13.3$ |
| s-Adiponectin (μg/ml) | 43 | $5.2 \pm 2.3$ | s-Adiponectin (μg/ml) | 25 | $10.0 \pm 4.4$ |
| s-Hs-CRP (mg/L) | 45 | $7.8 \pm 9.8$ | s-Hs-CRP (mg/L) | 25 | $2.9 \pm 5.3$ |

| *CSF measurement* |  |  | *CSF measurement* |  |  |
| --- | --- | --- | --- | --- | --- |
| CSF-Insulin (mU/L) | 45 | $0.33 \pm 0.12$ | CSF-Insulin (mU/L) | 17 | $1.5 \pm 1.6$ |
| CSF-Leptin (ng/ml) | 45 | $0.24 \pm 0.08$ | CSF-Leptin (ng/ml) | 17 | $0.20 \pm 0.10$ |

The pregnant subjects underwent elective cesarean section the morning after an overnight fast. A 10-ml venous blood sample was taken by venipuncture before infusion of Ringer-acetate solution. Before spinal anesthesia, an introducer needle was inserted into the interspinous ligament at L3-4, and a 25-gauge Whitacre needle or a 25-gauge Pajunk Pencil Point Spinal Needle was inserted through the introducer into the subarachnoid space. Ten milliliters of CSF were removed with a 10-ml syringe. Hemorrhagic samples were excluded. The first 0.5 ml of CSF was discarded. Study samples were transferred to polyethylene tubes, placed on ice, centrifuged, aliquoted, and stored (−80°C). Serum samples were similarly centrifuged, aliquoted, and stored. At the 5-year follow-up, women came to the laboratory after an over-night fast and CSF and blood samples were taken and processed in the same way as at the cesarean section. Dietary intake was assessed with a self-administered questionnaire for the three previous months. The questionnaire had a semi-quantitative food frequency design and was validated in Swedish men and non-pregnant women against a 4-day food record and assessments of 24-h energy expenditure and nitrogen excretion. From these comparisons, valid estimates of energy intake were obtained in normal weight, overweight, and obese subjects.(30)

### Biochemical analysis

Blood glucose and insulin were analysed with a Cobas Modular system (Roche Diagnostics, Risch, Switzerland) at the Clinical Chemistry Laboratory, Sahlgrenska University Hospital (accredited in accordance with the International Standard ISO 15189:2007). CSF insulin was analyzed with a double-antibody radioimmunoassay (Linco Research) at the Department of Clinical Science, Lund University. Leptin and adiponectin were measured with ELISA kits (R&D Systems) in the Clinical Neurochemistry Laboratory at Sahlgrenska University Hospital Mölndal. ELISA plates were read on a Vmax plate reader, and concentrations were determined with Softmax software (Molecular Devices). Insulin was measured in undiluted samples and adiponectin in 100-fold diluted samples. For leptin analysis, CSF samples were diluted 2-fold and serum samples 100-fold. HOMA-IR was calculated as (fasting glucose × fasting insulin)/22.5 and HOMA-B as (20 × fasting insulin)/(fasting glucose – 3.5).(31) GDF15 concentration was measured with Human GDF-15 Quantikine Elisa Kit (R&D Systems, Minneapolis, MN, USA). Serum GDF15 samples during and after pregnancy were diluted 1:64 and 1:4, respectively. CSF GDF15 samples were diluted 1:2. The intra- and inter-assay coefficients of variation (CVs) for GDF15 measurements were 1.7% and 7%, respectively.

### Statistical analysis

Variables are expressed as mean ± standard deviation (SD). Differences between pregnancy and follow-up were assessed using paired t-tests; between-group differences for fetal sex were assessed using independent t-tests and univariate tests. Associations were analyzed using Pearson correlations and linear regression models. Adjustments in linear regression models were made for gestational age, or for gestational age, maternal pre-pregnancy BMI, gestational weight gain and fetal sex. All tests were two-tailed and conducted at a 0.05 significance level.

## Results

### Pregnancy GDF15 levels were increased in both serum and CSF

During pregnancy, GDF15 was almost 200 times higher in serum and more than 3 times higher in CSF compared with levels five years after pregnancy (Table 2). The GDF15 ratio CSF:serum was, however, lower during pregnancy compared with follow up.

**Table 2.**
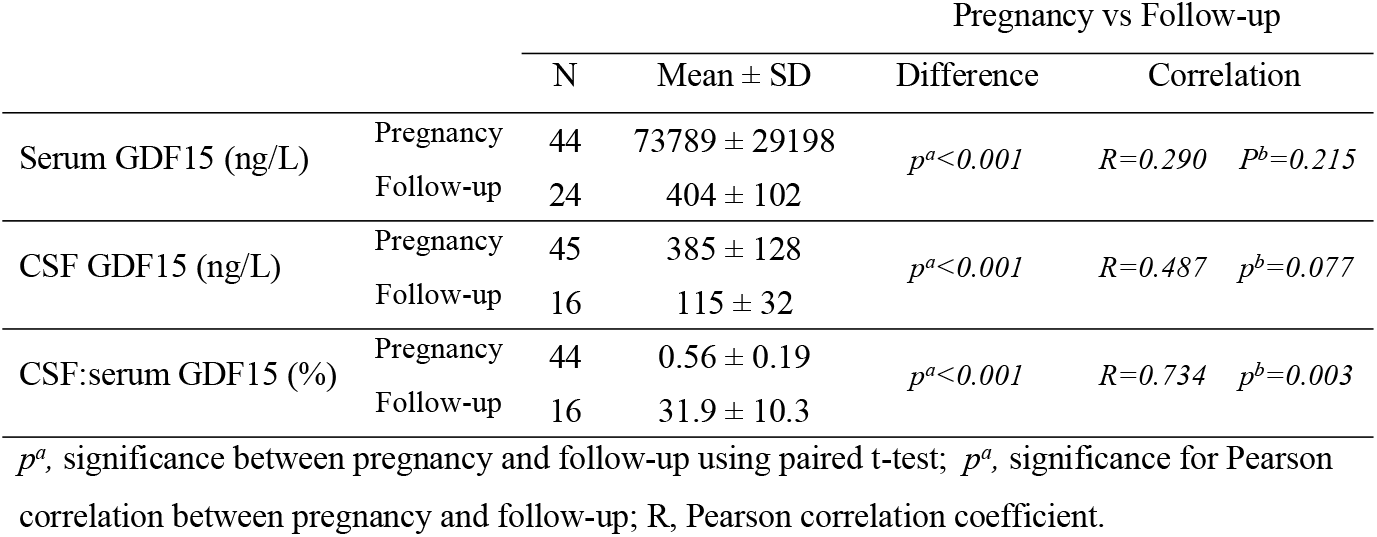
GDF15 in serum and CSF at delivery and at follow-up 5 years after delivery

Pregnancy GDF15 levels did not correlate with post-pregnancy GDF15 levels in either serum or CSF. However, a strong significant correlation was observed for the ratio CSF:serum between the two time points (Table 2).

CSF GDF15 was also compared with serum GDF15 within the two time points separately (Fig 1). During pregnancy, there was a clear and significant correlation (r=0.577; p<0.001) between GDF15 serum and CSF levels (Figure 1A). In contrast, 5 years after pregnancy there was no correlation between serum and CSF levels (Figure 1B).

**Figure 1.**
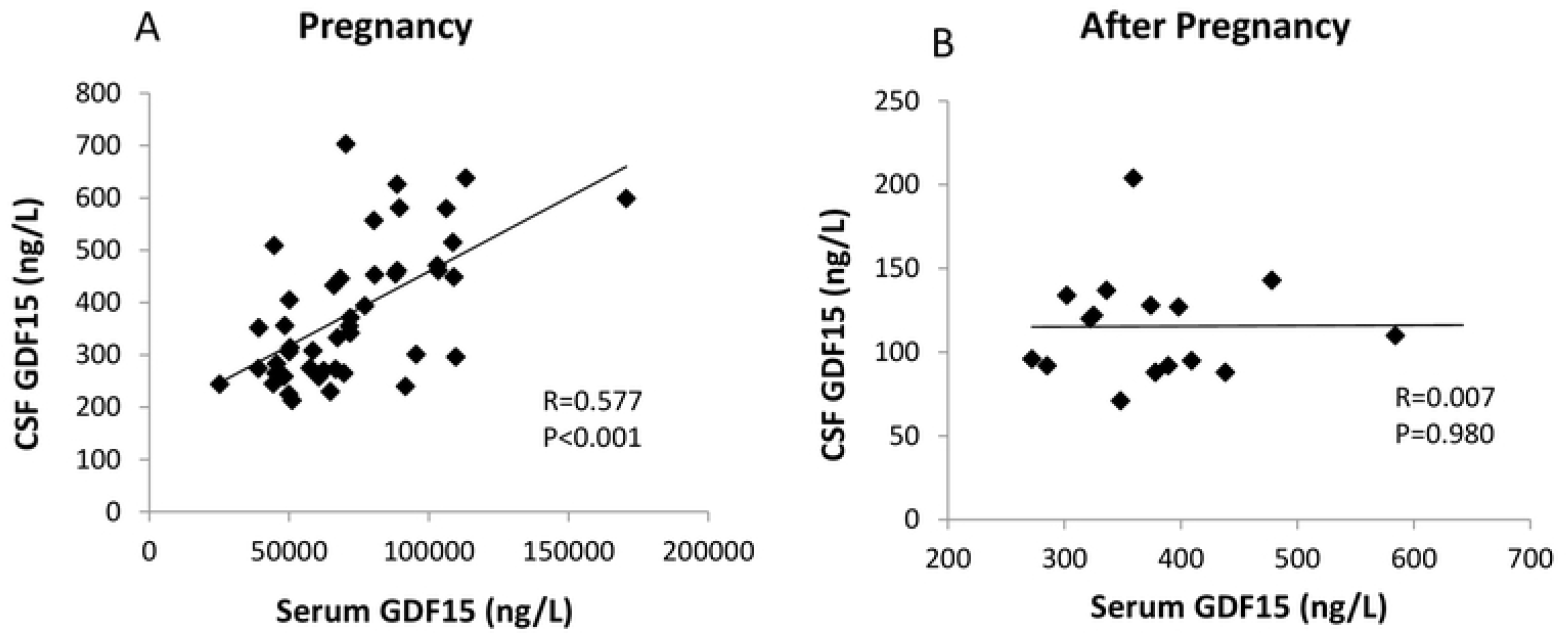
Correlations between GDF15 in cerebrospinal fluid and GDF15 in serum. Pearson correlation at A) cesarean section and B) 5 years after pregnancy.

### Women carrying a female fetus had the highest GDF15 levels in both serum and CSF

Women carrying a female fetus had significantly higher GDF15 levels both in serum and in CSF during pregnancy compared with women carrying a male fetus (Figure 2), whether or not the analysis was adjusted for gestational age, maternal age and BMI. There was no significant difference in the CSF:serum ratio of GDF15 between mothers carrying fetuses of different sex. After pregnancy, there were no differences in GDF15 in either serum or CSF between mothers that carried female vs. male fetuses (p > 0.6).

**Figure 2.**
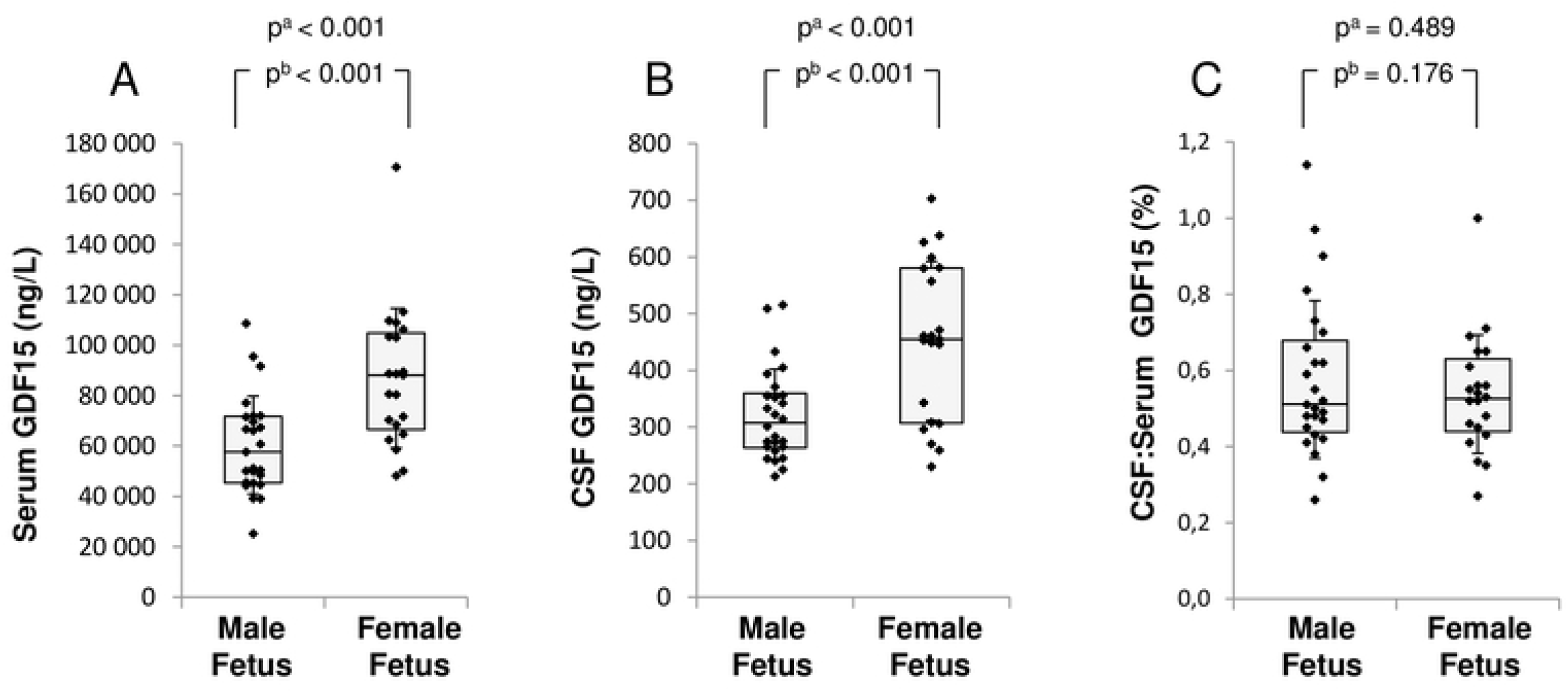
Maternal GDF15 levels at cesarean section depending on fetal sex. Box plot of GDF15 in A) serum, B) CSF or C) CSF:serum ratio. ^a^independent t-test; ^b^adjusted for maternal pre-pregnancy BMI, maternal age and gestational age. The box represents first quartile, median and third quartile; the whiskers represent standard deviation.

### Serum GDF15 associated with beta-cell function

At cesarean section, there were no associations between GDF15 in serum or in CSF with self-reported pre-pregnancy BMI (P=0.18) or gestational weight gain (P=0.92). There was a positive association for serum GDF15 with HOMA-B (Table 3), but no significant associations with glucose, insulin or HOMA-IR were found. Similarly, associations between serum GDF15 and the adipokines leptin and adiponectin, the inflammation marker hs-CRP, and birth weight were not significant. The association between GDF15 and placental weight was significant when adjusted for gestational age, but not in the fully adjusted model.

**Table 3.** Associations for serum and CSF GDF15 during pregnancy.

|  | Model A |  | Model B |  |
| --- | --- | --- | --- | --- |
| | $\beta$ | *p* | $\beta$ | *p* |
| *Serum GDF15* |  |  |  |  |
| p-Glucose | -0.181 | 0.269 | -0.286 | 0.165 |
| s-Insulin | -0.162 | 0.322 | -0.289 | 0.098 |
| HOMA-IR | -0.180 | 0.273 | -0.284 | 0.123 |
| HOMA-B | **0.449** | **0.003** | **0.590** | **0.001** |
| s-Leptin | -0.097 | 0.556 | -0.190 | 0.343 |
| s-Adiponectin | 0.234 | 0.155 | 0.239 | 0.277 |
| s-Hs-CRP | 0.220 | 0.154 | 0.106 | 0.564 |
| Placental weight | **0.325** | **0.044** | 0.289 | 0.191 |
| Birth weight | 0.100 | 0.543 | 0.019 | 0.932 |
| *CSF GDF15* |  |  |  |  |
| CSF-Insulin | **-0.318** | **0.038** | -0.232 | 0.275 |
| CSF-Leptin | -0.165 | 0.293 | -0.128 | 0.527 |

Model A- adjusted for gestational age at sampling
Model B – adjusted for gestational age, maternal age, maternal pre-pregnancy BMI, gestational weight gain and fetal sex.

There was a negative association between CSF GDF15 and insulin when adjusted for gestational age, but not in the fully adjusted model (Table 3). There was no association with CSF leptin.

At follow up, none of the blood or CSF measurements correlated significantly with GDF15 (data not shown). There were no significant correlations between GDF15 and energy or macronutrient intake at delivery or at follow-up.

## Discussion

For the first time, GDF15 levels were measured in CSF of pregnant women. We showed increased levels of GDF15 in both CSF and serum compared with the non-pregnant state, whereas the ratio CSF:serum was lower during pregnancy. Furthermore, women carrying a female fetus had higher GDF15 levels in both serum and CSF.

We have previously shown prospectively in a different cohort that serum GDF15 was increased up to 200 times in trimester 3 compared with the non-pregnant state.(12) We measured similar elevated serum levels in the current population, but also showed an increase in GDF15 in the CSF during pregnancy. The (approximately 3-fold) increase in CSF was smaller than observed in serum, thus the ratio of CSF:serum was decreased during pregnancy. GDF15 has been proposed to regulate food intake, to be involved in taste aversion and possibly in food choice, as well as in improvement of glucose tolerance.(3–5, 32, 33) High serum GDF15 levels during pregnancy have previously also been linked to nausea and hyperemesis gravidarum, and in our previous study we showed an association with beta-cell function as measured by HOMA-B.(12, 16–18) However, anorexia and cachexia, which would be expected at high GDF15 levels, does not normally occur during pregnancy. Pregnant women increase food intake and gain a substantial amount of weight (for example, normal-weight women in our prospective cohort gained 10.5 kg body weight of which 4 kg was fat mass(34)). How these greatly increased GDF15 levels coexist with increased energy intake during pregnancy is not yet known. Although the main effects of GDF15 are believed to be regulated through the GFRAL receptor of the brainstem,(1, 2, 22, 35) GFRAL-independent effects centrally or peripherally cannot be excluded (36). GFRAL has been shown to be exclusively expressed in the AP and NTS regions of the brainstem in the non-pregnant state,(1, 2, 22) although no studies have examined maternal GFRAL expression during pregnancy.

AP has a permeable BBB, which means that the GFRAL receptor is exposed to circulating ligands from both blood and CSF, whereas NTS is situated behind a less permeable BBB.(24) In murine models, both peripheral and central injections of GDF15 reduce food intake and cause weight loss.(22, 25) Effects of increased GDF15 in CSF on human physiology are not known. GDF15 in CSF has been measured only in a few studies, never in pregnancy, but in conjunction with its potential role in CNS disease. In these studies GDF15 was shown to be increased in CSF of patients with neurodegenerative disorders such as multiple sclerosis(26) and Parkinson’s disease(27) or in glioblastoma patients(28). Patients in these studies typically had GDF15 levels of between 200-300 pg/ml in CSF (approximately 50% higher than matching controls), i.e. a smaller increase than we found in pregnancy in the present study. The control subjects in previous studies showed similar CSF levels to our non-pregnant subjects at follow-up.(26, 27) Unfortunately, none of the previous studies with CSF levels of GDF15 performed parallel measurements of serum GDF15 so comparisons of CSF:serum ratios are not available.

In the non-pregnant state, we saw no correlation between GDF15 in CSF and serum, which may indicate that a proportion of GDF15 in CSF is produced within the CNS. In pregnancy, however, a state with large GDF15 increases, CSF levels correlated with serum levels. One explanation for this could be that large peripheral increases in GDF15 from placental expression,(10) leads to increased transport (active or passive) from blood to CSF across the BBB. Interestingly, even though there was no association found with either CSF or serum GDF15 levels when comparing the pregnant vs. non-pregnant state, the CSF:serum GDF15 ratio was strongly correlated between states. This could be interpreted as women with a high degree of GDF15 transport across the BBB in the non-pregnant state may also have a high degree of transport in the pregnant state. However, as mentioned above, it should be noted that all GDF15 in CSF is presumably not from a peripheral origin. Expression of GDF15 in CNS and release into the CSF has been shown in murine studies,(37) and CNS expression has been documented in the open-access Brain Atlas resource of the Human Protein Atlas.(38)

We have previously shown, in a different cohort of pregnant women, higher GDF15 serum levels in women carrying a female compared with male fetus.(12) We confirmed that finding in the current cohort, and also showed that CSF levels of GDF15 were higher in the women carrying a female fetus. Since this is a new observation, the mechanism for higher GDF15 levels in female pregnancies is not known. However, sexual dimorphism is found for placental transcription of other endocrine molecules found at high levels during pregnancy (such as human chorionic gonadotropin, HCG), where sex chromosomes has been suggested to play an important role.(39) Even though the reason for increased GDF15 levels is not known, one could speculate that the higher degree of nausea found in women carrying girls (40–43) might be linked to the higher GDF15 concentrations in serum and/or CSF. There was no difference in CSF:serum ratio of GDF15 for women with male versus female fetuses, which is as expected if CSF levels are mainly determined by peripheral levels.

We also confirmed in this new cohort of women our previously shown association between serum GDF15 and HOMA-B in serum. Additionally, we observed a negative association of CSF GDF15 with CSF insulin, but no association with CSF leptin. Both insulin and leptin are believed to be involved in the central regulation of energy balance and peripheral glucose metabolism during pregnancy.(44) Both hormones stimulate POMC and inhibit AgRP/NPY neuronal activity and are therefore implicated in decreased food intake. In the present study, we found no change in CSF leptin during pregnancy compared to follow-up, whereas CSF insulin was lower during pregnancy compared to follow-up (*p*=0.01). In agreement with our previous study of serum GDF15 in pregnancy, we did not see a correlation with gestational weight gain. We also did not observe any association with self-reported dietary intake (data not shown), although it should be noted that only approximately half of the women returned a completed dietary questionnaire.

As most research on GDF15 in the CNS has been performed in animal studies, this study adds important new knowledge in the human. The design of the study with women acting as their own controls is a strength of the study. However, limitations include the shortfall in number of women attending the follow-up CSF sampling. This reduces power and could open for potential bias. The women attending both visits had lower BMI, glucose and insulin compared with the women only sampled at pregnancy. Importantly, however, there were no differences in GDF15 levels, and with only performing paired longitudinal analysis for changes between the two time points the bias should not influence the results of the present study. Also, we did not have information about pregnancy induced nausea, which would have been valuable to evaluate the role played by CSF and serum GDF15 in this respect.

In conclusion, we have measured GDF15 in CSF of pregnant women and compared the levels longitudinally to the non-pregnant state. We showed increased levels of GDF15 in both CSF and serum during pregnancy compared with follow-up, and that both CSF and serum levels were highest in women carrying a female fetus. We propose that effects of GDF15 during pregnancy can be mediated by changes in both CSF and serum levels. These types of human studies are important to start elucidating how GDF15 might be centrally regulated in its proposed actions involving appetite, nausea and glucose metabolism.

## Acknowledgements

We would like to thank anesthetists Aurimantas Pelanis and Ove Karlsson for CSF sampling, and Ulf Andreasson for help with biochemical analysis of the original study

## Data availability statement

The data that support the findings of this study are available from the corresponding author upon reasonable request.

